# A novel cellular structure in the photoreceptors of insectivorous birds

**DOI:** 10.1101/551945

**Authors:** Luke P. Tyrrell, Leandro B.C. Teixeira, Richard R. Dubielzig, Diana Pita, Patrice Baumhardt, Bret A. Moore, Esteban Fernández-Juricic

## Abstract

The keen visual systems of birds have been relatively well-studied. The foundations of avian vision rest on their cone and rod photoreceptors. Most birds use four cone photoreceptor types for color vision, a fifth cone for achromatic tasks, and a rod for low-light levels. The cones, along with their oil droplets, and rods are conserved across birds – with the exception of a few shifts in spectral sensitivity – despite taxonomic, behavioral and ecological differences. Here, however, we describe a novel photoreceptor in a group of New World flycatchers (*Empidonax* spp.) in which the traditional oil droplet is replaced with a complex of electron-dense megamitochondria surrounded by hundreds of small, orange oil droplets. These photoreceptors were unevenly distributed across the retina, being present in the central region (including in the fovea), but absent from the retinal periphery and the *area temporalis*. Many bird species have had their oil droplets and photoreceptors characterized, but only the two flycatchers described here (*E. virescens and E. minimus*) possess this unusual structure. We discuss the potential functional significance of the unique sub-cellular structure in these photoreceptors providing an additional visual channel for these small predatory songbirds.

## Introduction

Birds are widely recognized for their specialized visual abilities^1^, which enable them to engage in many diverse modes of life. For example, birds can be diurnal or nocturnal, predators and/or prey, and flighted or flightless. The capabilities of avian vision have led to much interest in the evolutionary design of avian visual systems^2^. The foundation of avian vision lies in their six photoreceptor types^3^, compared for example to just four photoreceptors in humans. Birds, from highly active predators such as hawks and falcons to passive foragers such as emus and sparrows, have four single cone photoreceptors for color vision (ultraviolet or violet, short-wavelength, medium-wavelength, and long-wavelength sensitive), a double cone for achromatic discrimination/motion vision, and a rod for vision in dim light^3,4^. Within each cone photoreceptor, birds have carotenoid-filled organelles called oil droplets, which are involved in filtering light before it reaches the visual pigments, enhancing color discrimination^3^. Bird species differ in the distribution, abundance, and absorbance spectra of these six photoreceptors^3^, but the six fundamental building blocks are the same across species.

In this study, we report the discovery of a potentially seventh photoreceptor that contains a cellular organelle not found in this form in any other vertebrate retina with important functional consequences for visual perception. We found this photoreceptor in two species of New World flycatchers of the genus *Empidonax* (*E. virescens and E. minimus*). We describe this new cellular structure using light microscopy and transmission electron microscopy, analyze its spectral composition using microspectrophotometry, and establish the distribution and densities of these novel cones relative to the traditional cones across the retina. Additionally, we displayed the presence/absence of this cellular structure in the avian phylogeny based on species whose cones and oil droplets have been previously characterized. Overall, this cellular structure may allow these sit-and-wait flycatchers to see their world from a different perspective than other animals.

## Results

We found that *Empidonax* flycatchers, like all birds, had four single cone photoreceptors that each contained a spherical oil droplet in the inner segment of the photoreceptor. Like other birds, the principal member of the *Empidonax* double cone also contained a spherical oil droplet. Each type of cone had a different colored oil droplet that could be readily visualized under simple light microscopy (Fig. 1A). In addition to these five traditional cones and their corresponding oil droplets, we found that the *Empidonax* flycatcher retina contained what is probably an additional photoreceptor with a novel, orange, conical structure in the apical end of the inner segment (Fig. 1B).

**Fig. 1.**
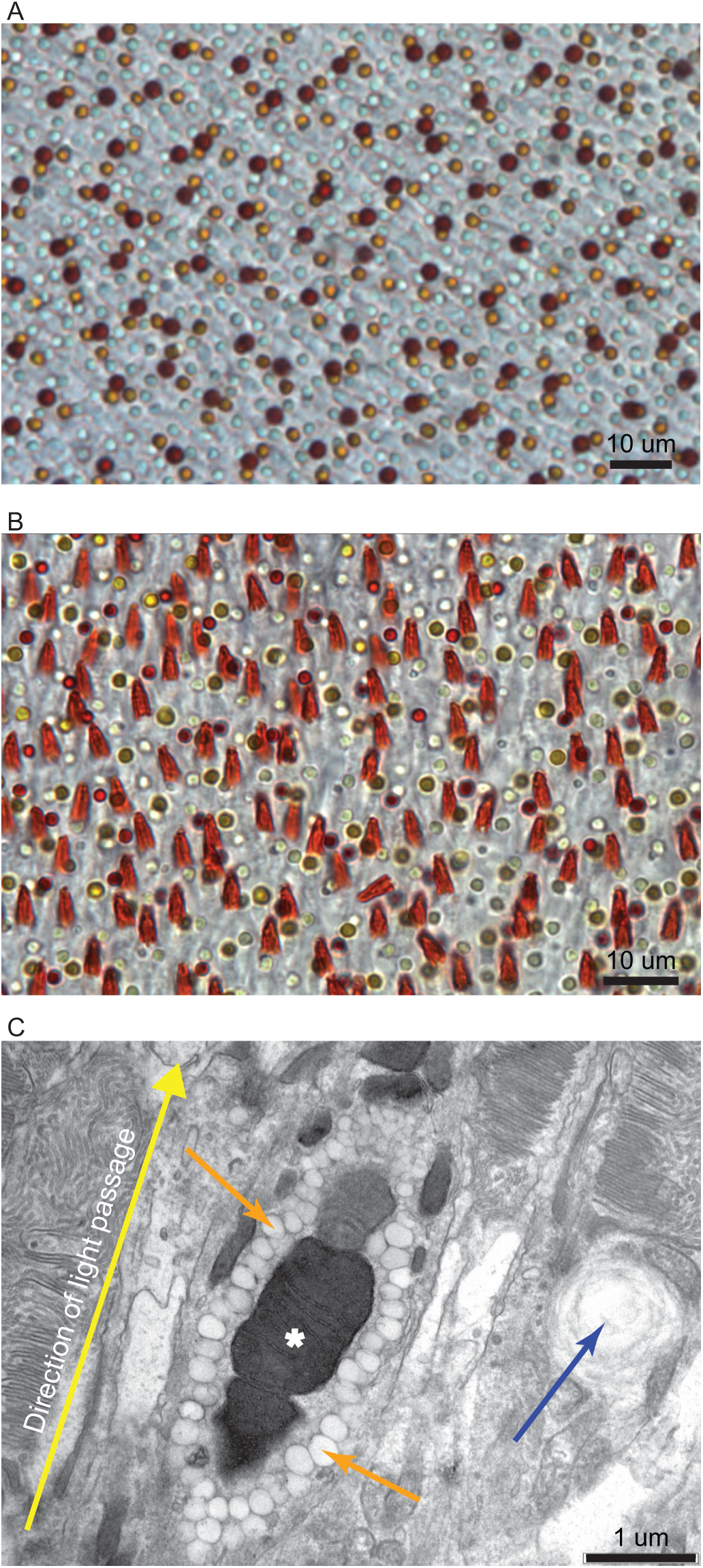
Light and electron microscopy of oil droplets and MMOD-complexes. (**A**) Image of a white-throated sparrow (*Zonotrichia albicollis*) retina wholemount under light microscopy showing the five traditional oil droplets types. (**B**) Image of an Acadian flycatcher (*Empidonax virescens*) retina wholemount under light microscopy showing the five traditional oil droplet types and the additional orange, conical structures belonging to the new photoreceptor. (**C**) Transmission electron microscopy image of the orange, conical structures reveals that they are electron-dense megamitochondria surrounded by many small oil droplets. The white asterisk denotes the megamitochondria, the orange arrows denote the small oil droplets that provide the orange coloration seen in panel **B**, and the blue arrow indicates a traditional oil droplet from the neighboring photoreceptor. The retinae in panels **A** and **B** have not been stained or artificially labeled in any way.

### Electron Microscopy

Transmission electron microscopy revealed that this novel structure consisted of a cluster of electron-dense giant mitochondria (hereafter megamitochondria) that were surrounded by hundreds of small oil droplets (Fig. 1C, Fig. S1). These small oil droplets were the likely source of the orange coloration seen under light microscopy. These megamitochondria were over two times the diameter of the mitochondria in surrounding photoreceptors and the small oil droplets associated with it had a diameter over seven times smaller than the traditional oil droplets found in the other cone types (Fig. 1C). Given the unusual combination of megamitochondria and small oil droplets, we coined the term megamitochondria-small oil droplet complex (hereafter, MMOD-complex) to refer to this new cell structure.

The MMOD-complexes identified using electron microscopy were of the same shape and size as those first observed on retinal wholemounts (Fig. 1, Fig. S1). Additionally, the MMOD-complexes observed using electron microscopy were only present at the center of the flycatcher retinae. No similar structures were present in the retinal periphery of flycatchers or in any part of the retinae from two other species that we used for comparison purposes under electron microscopy (white-throated sparrow and house sparrow, see Methods).

We also observed connecting cilia that linked the MMOD-complex to a photoreceptor outer segment in flycatchers (Fig. S1). Therefore, we determined that the MMOD-complex was likely housed at the apical end of the inner segment of a photoreceptor, just distal to the traditional mitochondrial-rich ellipsoid. This is where a traditional oil droplet would normally be located, which is absent in this new photoreceptor type. In addition to the connecting cilia between the MMOD-complex and the photoreceptor outer segment, it would seem logical that any structure occupying space in the photoreceptor array of the retina would serve a photoreceptive function. Diurnal birds have retinas with dense photoreceptor arrays that have little or no gap between adjacent photoreceptors. Non-photoreceptive cells inserted into the photoreceptor array occupy valuable ‘real estate’ and would reduce visual acuity, seemingly unnecessarily, although exceptions to this line of thinking are possible. Furthermore, it is not likely that the MMOD-complex serves a specialized energy source for other photoreceptors or other cells in the retina because cellular energy that is produced in a cell, generally, must remain in the cell that produced it and cannot be shared with neighboring cells^5^. These lines of thought, along with the connecting cilia between the MMOD-complex and a photoreceptive outer segment, are why we think the most likely scenario is that the MMOD-complex is part of a photoreceptor.

### Microspectrophotometry

We found that the MMOD-complexes had λ_mid_ = 565 nm, which corroborates their orange coloration under bright microscopy (Fig. 2b). Their λ_mid_ falls between the yellow Y-type oil droplets (516 nm) and the red R-type oil droplets (580 nm). The small oil droplets surrounding the megamitochondria work as long-pass filters, letting light longer the 565 nm pass through and absorbing shorter wavelengths. Full microspectrophotometry results can be found in Table S1.

**Fig. 2.**
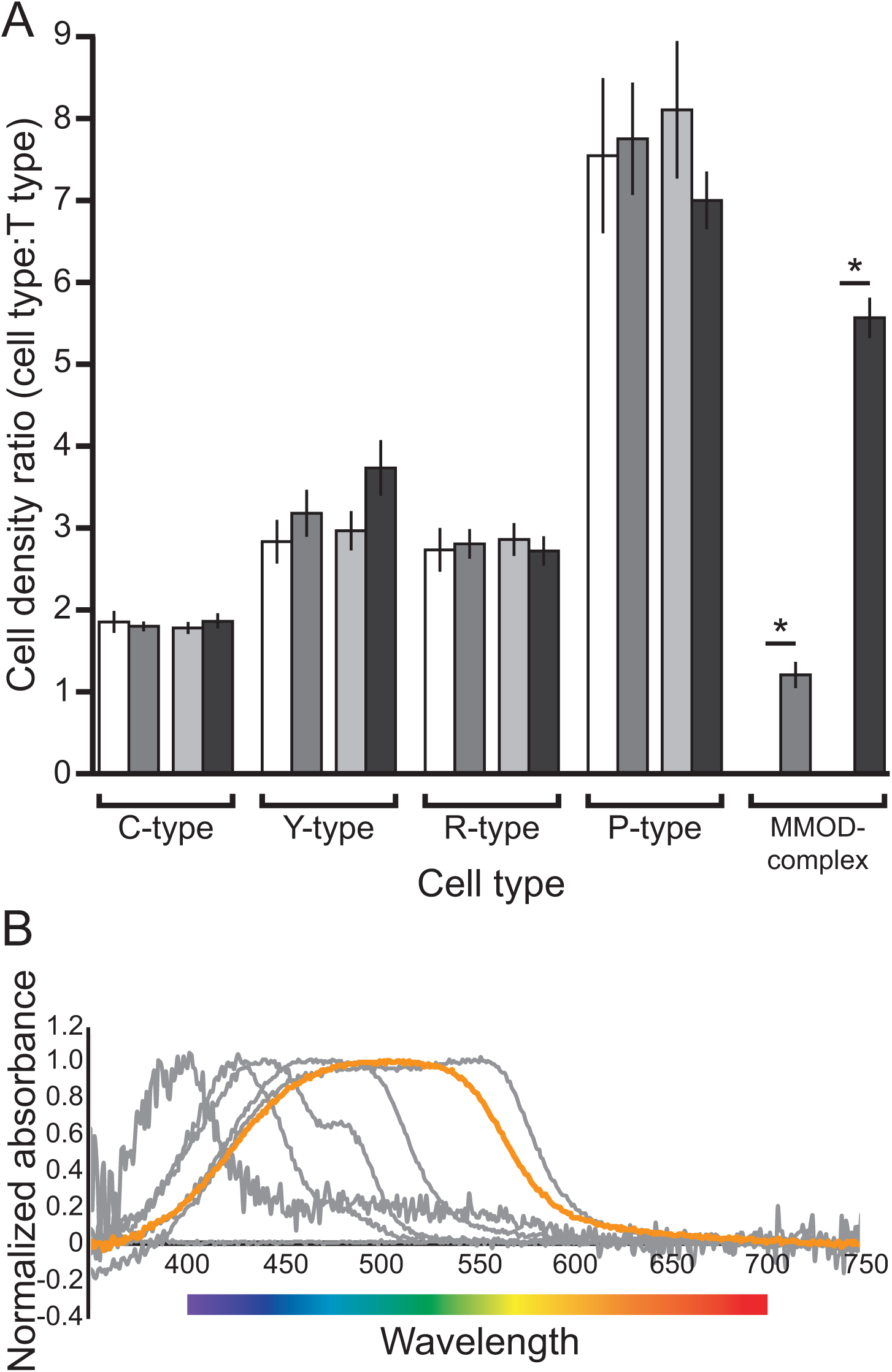
Relative densities and absorbance spectra of *Empidonax* oil droplets. (**A**) Ratios of oil droplet density in the retina of typical passerines^3^ (white bars), the entire retina of *Empidonax* flycatchers (medium gray bars), areas of *Empidonax* retinae where MMOD-complex photoreceptors are absent (light gray bars), and areas of *Empidonax* retinae where MMOD-complex photoreceptors are present (dark gray bars). Values on the Y-axis are the ratios of the different oil droplet types on the X-axis to T-type oil droplets. Statistically significant differences are denoted by an asterisk. (**B**) Absorbance spectra of *Empidonax* oil droplets. The orange line corresponds to the MMOD-complex. The gray lines correspond to the traditional oil droplets.

### Photoreceptor Distribution

The *Empidonax* flycatcher retinae we examined contained a centrally placed fovea (i.e., an invagination in the retinal tissue that corresponds to the highest photoreceptor densities in the retina) and an *area temporalis* (i.e., an increase in cell density without a foveal pit in the temporal region of the retina) (Fig. 4). The peak photoreceptor density for all counted cell types (all single cones, double cones and MMOD-complex photoreceptors) was 118,300 ± 9,022 cells/mm^2^ and 58,533 ± 4,791 cells/mm^2^ in the fovea and *area temporalis*, respectively (Fig. 3a). The peak foveal density was 33,700 ± 3,649 cells/mm^2^ and 85,600 ± 5,757 cells/mm^2^ for MMOD-complex photoreceptors (Fig. 3i) and traditional oil droplets (Fig. 3b), respectively. Interestingly, the MMOD-complex photoreceptors followed a different distribution pattern than the other photoreceptors. Traditional cones were present throughout the retina and their density decreased moving away from the fovea towards the periphery. However, MMOD-complex photoreceptors were only present in the central region of the retina, including in the fovea. Their density decreased slightly moving away from the fovea, but then these MMOD-complex photoreceptors suddenly disappeared altogether (Fig. 3i-j). There were no MMOD-complex photoreceptors present in the retinal periphery or in the *area temporalis*. Therefore, the MMOD-complex region of the retina primarily subtends the lateral and horizontal section of the visual field (i.e., to the sides of the head without a downward or upward bias towards the ground or sky). The peak foveal density of T-type, C-type, Y-type, R-type, and P-type oil droplets was 6300 ± 1258 cells/mm^2^, 9700 ± 551 cells/mm^2^, 23600 ± 1720 cells/mm^2^, 13400 ± 683 cells/mm^2^, and 36100 ± 3515 cells/mm^2^, respectively (Fig. 3d-h).

**Fig. 3.**
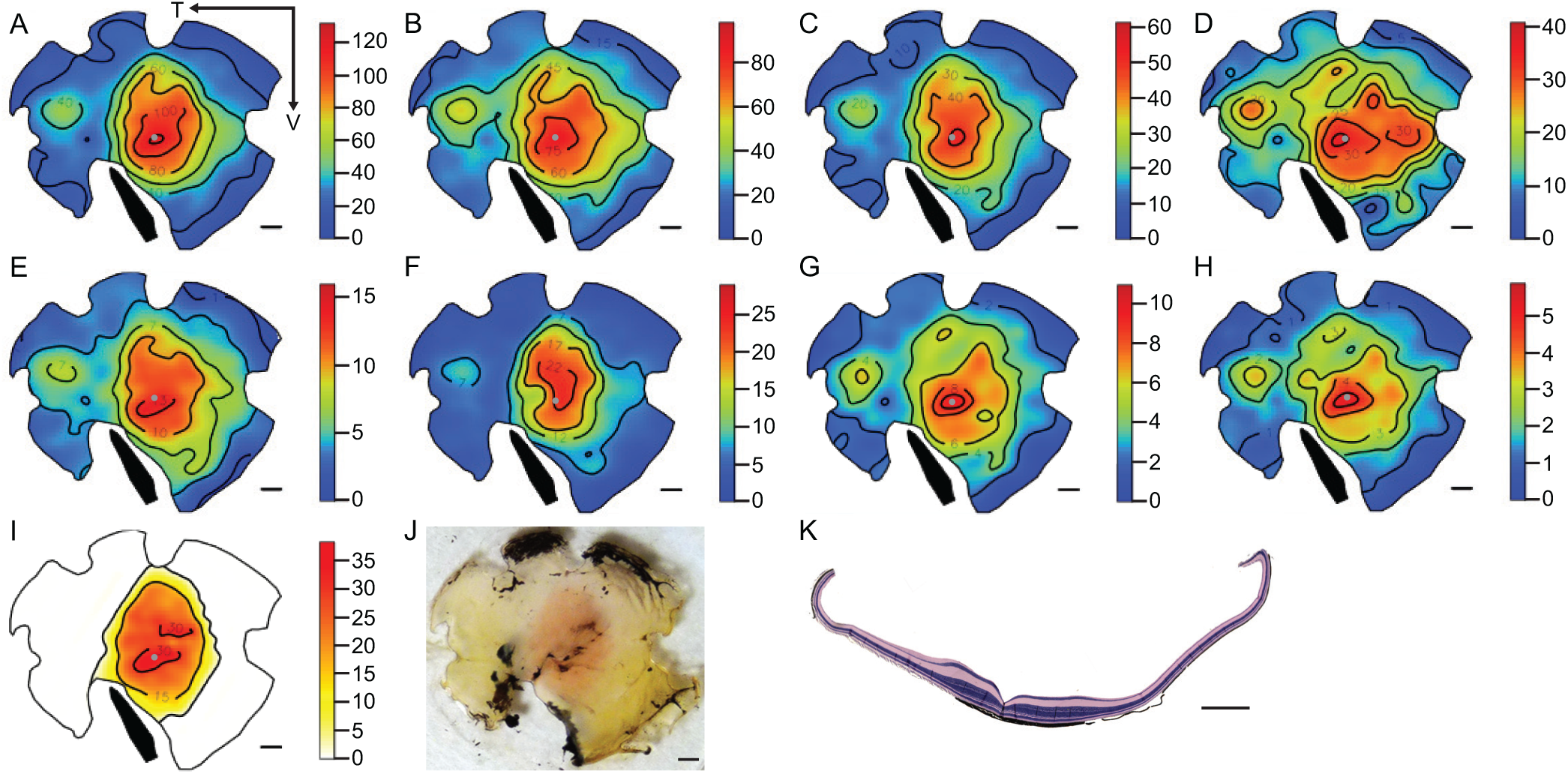
Retinal topography and histology of *Empidonax* flycatchers. (**A**) All cones, (**B**) all traditional cones, (**C**) all single cones, (**D**) P-type oil droplets, (**E**) R-type oil droplets, (**F**) Y-type oil droplets, (**G**) C-type oil droplets, (**H**) T-type oil droplets, (**I**) MMOD-complex photoreceptors, (**J**) trans-illuminated photograph of the retinal wholemount with the orange central area of MMOD-complex photoreceptors clearly visible, (**K**) cross section showing the fovea (invagination in the retinal tissue) and retinal thickening corresponding to the MMOD-complex photoreceptor region. All numbers are X x10^3^, all scale bars are 1 mm, the arrows in panel A indicate the temporal (T) and ventral (V) directions, gray dots indicate the position of the fovea, and black bands indicate the position of the pecten.

**Fig. 4.**
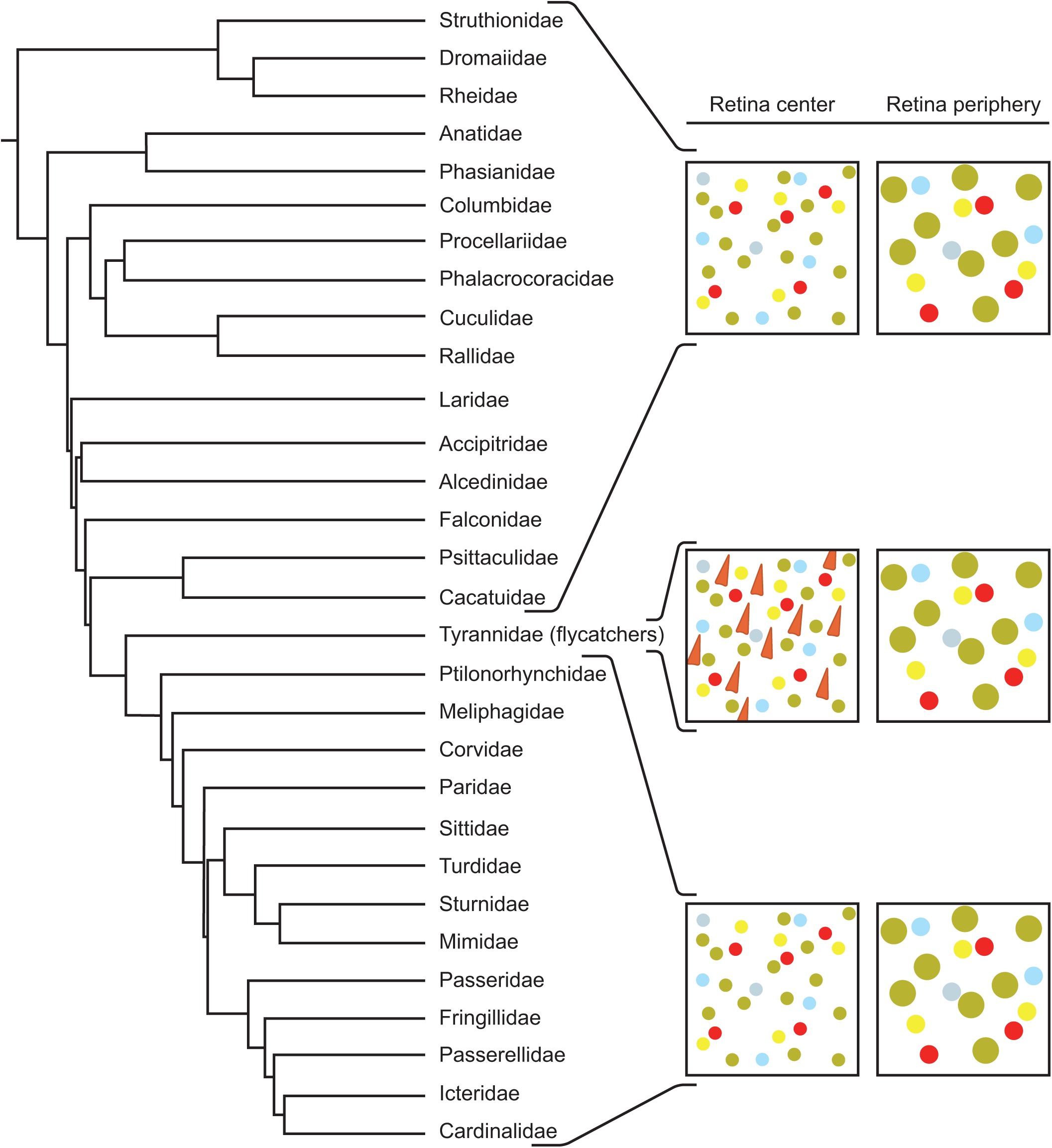
Photoreceptors across the avian phylogeny. Across many studied avian Families, only the New World flycatchers of Family Tyrannidae show deviation from the typical avian photoreceptor classes.

We found no significant differences between typical passerine retinae and *Empidonax* retinae regarding relative ratios of C-type, Y-type, R-type, or P-type oil droplets (Fig. 2a, Table S2). Unsurprisingly, there was a significant difference in the relative abundance of MMOD-complex photoreceptors between typical passerine retinae – which lack the MMOD-complex – and *Empidonax* retinae (Fig. 2a, Table S2).

When comparing the center region of *Empidonax* retinae that contained MMOD-complex cells to the peripheral region of *Empidonax* retinae that lacked MMOD-complex cells, we found no significant differences for C-type, Y-type, R-type, or P-type oil droplet ratios (Fig. 2a, Table S2). This means that no specific cone type is being disproportionately replaced by MMOD-complex cells. Again, there was a significant difference in the relative density of the MMOD-complex cells between counting locations that lacked MMOD-complex cells – peripheral region – and counting locations that contained MMOD-complex cells – central region (Fig. 2a, Table S2).

### Cross Sections

The MMOD-complex region covered the central 27.4 ± 1.7 mm^2^ of the retina (or 24.6 ± 0.9% of the whole retina). The diameter of the retinal region containing MMOD-complexes measured on the naso-temporal axis of wholemounts (4.44 ± 0.14 mm) was similar to the naso-temporal diameter of the thickened retinal region on the cross sections (4.24 ± 0.37 mm). This suggests that the MMOD-complex photoreceptors may increase the demand for cells in other retinal layers, leading to a substantial thickening of the retinal tissue in regions where MMOD-complexes are present.

### Phylogenetic Representation

We reviewed the literature and also used unpublished data describing the retinal photoreceptor layer (i.e., wholemounts of fresh retinas where the oil droplets were visible under bright and/or florescent microscopy) of different avian species (Table S3). Considering the 48 bird species for which information was available, the MMOD-complex was only found in the two flycatchers described here (*E. virescens and E. minimus*) belonging to the Family Tyrannidae, Order Passeriformes (Fig. 4). However, the MMOD-complex is absent in the other 46 species that belong to other taxanomic Families or Orders (Fig. 4).

## Discussion

The MMOD-complex is different in a number of ways from similar retinal structures that have been described in other organisms. Birds from many taxa have been studied previously and all share the same classes of photoreceptors, and none showed the MMOD-complex found in the two New World flycatchers reported here (Fig. 4). Pigeons (*Columba livia*) do have red microdroplets in a specialized area of the retina called the “red spot” or “red field” that is well positioned for viewing green backgrounds in the pigeons’ binocular field^6,7^. Unlike flycatchers, however, the pigeons’ microdroplets are present in addition to a principal oil droplet within the same inner segment and are not associated with a mitochondria^6^. Additionally, lamprey photoreceptors contain either yellow/orange pigments (but not droplets) or large, electron-dense mitochondria. However, the pigments and electron-dense mitochondria have not been found in the same photoreceptor^8^, as is the case with flycatchers.

Megamitochondria have been found in the retina of tree shrews (*Tupaia* spp.)^9^ and zebrafish (*Danio rerio*)^10^. However, the morphological novelty of the flycatcher megamitochondria is that they are part of a larger complex that includes hundreds of small oil droplets. These small oil droplets surround the megamitochondria, absorb visible light below 565 nm (λ_mid_), and allow longer wavelengths – oranges and reds – to pass through (Fig. 2B, Table S1). Therefore, the small oil droplets flood the megamitochondria (and the photoreceptor outer segment) with long wavelength light. Put another way, the small oil droplets prevent short and medium wavelength light from reaching the megamitochondria, thereby isolating any effects of long wavelength light.

The cones of vertebrates have organelles in their inner segments called ellipsoids, which are accumulations of mitochondria^11,12^. These ellipsoids are even present in vertebrates with oil droplets, like birds^13^. From a developmental perspective, it is possible that the MMOD-complex is a modification of these ellipsoids because in some species of fish without oil droplets, the ellipsoids appear to resemble spatially and functionally the oil droplets^14^ and the ellipsoids have been proposed to give rise to cone droplets through a process of metabolic turnover^15^.

Interestingly, the MMOD-complex photoreceptors do not follow the same distribution pattern across the retina as other photoreceptors do. Traditionally, cones are at very high density in the *fovea centralis* and decrease in density towards the retinal periphery^16^, which is also the general case for traditional cones in the flycatcher retina (Fig. 3A-H). This heterogeneous distribution is because (1) cones are physiologically expensive^17^, and (2) the brain cannot simultaneously devote visual attention to every photoreceptor^17^. However, the MMOD-complex photoreceptors are only present at the center of the retina, with their highest density around the *fovea centralis*, decreasing away from the *fovea centralis* but abruptly disappearing from the retinal periphery entirely. In total, the MMOD-complex photoreceptors are only present in the central ∼25% of the retinal surface area (Fig. 3I-J, Fig. S2). The uncommon distribution pattern of MMOD-complexes appears to be an exaggeration of the aforementioned traditional cone distribution pattern. Therefore, we speculate that the MMOD-complex photoreceptors may be even more physiologically-costly than traditional cones, which limits their presence exclusively to the region where they are most beneficial for the animal. In addition to the *fovea centralis*, flycatchers also have an *area temporalis* in the temporal region of the retina that has elevated densities of traditional cones (Fig. 3A-H). But even the *area temporalis* is devoid of MMOD-complexes (Fig. 3I).

The MMOD-complex appears to be present in addition to the traditional oil droplet types, rather than at the expense of any one particular type. All five traditional oil droplet types are present throughout the entire flycatcher retina in similar ratios to what one would expect to find in any other passerine songbird (Fig. 2A, Table S2)^3^. Additionally, there are no significant differences between the ratios of traditional oil droplets in regions of the retina with and without MMOD-complex photoreceptors (Fig. 2A, Table S2). Furthermore, the extra photoreceptor density in the central region of the retina (i.e., where the MMOD-complex photoreceptors are present) is matched by an area of increased retinal ganglion cell density^18^ and a substantial thickening of the retinal tissue (Fig. 3K, Fig. S2). Both lines of evidence suggest that there is a need for more neurons in layers of the retina downstream from the MMOD-complex photoreceptors and support the idea that the MMOD-complex photoreceptors may serve a separate rather than incremental function relative to the other photoreceptors.

The function of similar structures in other organisms are not entirely understood, and it is possible that the MMOD-complex could be serving any number of functions simultaneously. Establishing the definitive function of the MMOD-complex was beyond the scope of this study. To provide some context, however, we can make some non-exhaustive speculations about its function. For example, zebrafish rods with experimentally expressed megamitochondria produce over twice as much ATP as wild-type rods that lack megamitochondria^10^. Generally, ATP produced within a cell must be used by that cell rather than sharing energy with the entire retina^5^. Therefore, megamitochondria are likely energy powerhouses for only the photoreceptor that contains them. The small orange oil droplets of the MMOD-complex could further increase megamitochondrial energy production. Cytochrome *c* oxidase – the terminal enzyme in the electron transport chain that is responsible for energy production in mitochondria – is a photoacceptor that increases its activity and energy production under long-wavelength visible light^19–22^. White light – which is the presence of all visible wavelengths – down-regulates other enzymes in the electron transport chain that are photoacceptors of shorter wavelengths^19^. Therefore, white light results in no net change of energy production, whereas long-wavelength light increases overall energy production^19^. Thus, by absorbing short-wavelengths, the flycatchers’ orange oil droplets only transmit the energizing long-wavelengths onto the megamitochondria. Filtering out short-wavelengths would also protect the retina from photo-oxidative damage^8,23,24^. Considering the high oxidative stress already present from extra energy production, these protective aspects of MMOD-complex could reduce the probability of apoptosis^24,25^.

With potentially large quantities of energy available to the flycatchers’ MMOD-complex photoreceptor, it is likely capable of more rapid response rates^26–28^. Rapid photoreceptor response rates may yield higher temporal visual resolution and spatio-temporal tracking of motion^29–31^, which would ultimately give flycatchers a photoreceptor potentially specialized in motion detection and motion tracking. This specialization can be particularly useful for flycatchers because they use a sit-and-wait hunting strategy^32^ that requires considerable motion-tracking as individuals sit almost motionless while their flying prey (flies, mosquitos, etc.) move at high speeds^33^. Therefore, flycatchers must detect and track the absolute velocity of their prey rather than the slower, relative velocity required by predators that actively chase prey (e.g., swifts and swallows). *Empidonax* flycatchers often hunt against blue or green backgrounds. The filtering effect of the orange oil droplets may even shield the potentially motion-sensitive MMOD-complex photoreceptors from the motion of green leaves blowing in the wind, isolating the motion of the insect.

In summary, the retina of flycatchers from a single Family of songbirds have evolved a novel cellular structure in what is probably a photoreceptor that may confer some benefits in terms of detecting, tracking, and capturing fast-moving prey. Potential future approaches to establish the function of this structure should include visual pigment absorbance spectra to establish the sensitivity of the visual pigment of the photoreceptor with the MMOD-complex, wavelength-specific electroretinograms to determine the critical flicker fusion frequency of MMOD-complex photoreceptors compared to the other photoreceptor types, optokinetic response to quantify motion tracking ability, retinal oximetry to determine the energetic demands of these photoreceptors, and behavioral experiments to assess the ability of flycatchers to catch prey under different wavelength specific light conditions.

## Methods

### Animals

Three adult Acadian flycatchers (*Empidonax virescens*), one adult least flycatcher (*E. minimus*), one adult white-throated sparrow (*Zonotrichia albicollis*), and one adult house sparrow (*Passer domesticus*) were captured using mist nets in Tippecanoe County, Indiana. Animals were euthanized by CO_2_-asphyxiation on the same day of capture, then immediately processed for retinal tissue analysis or preservation. All experimentas were approved by and carried out in accordance with the regulations of The Purdue Institutional Animal Care and Use Committee (protocols 1201000567 and 1112000398).

### Electron Microscopy

To determine the subcellular structure of the novel structures, the right eyes of one Acadian flycatcher and one house sparrow were examined using transmission electron microscopy (TEM). After enucleation, hemisection, and vitreous humor extraction, the eyes were fixed in 2.5% glutaraldehyde in 0.1 M phosphate buffered saline (pH 7.4) at 4°C overnight. 2 mm^2^ sections of the house sparrow retina (three separate sections), the periphery of the Acadian flycatcher retina (one section), and the center of the Acadian flycatcher retina (two separate sections) were trimmed and embedded in epon resin. The house sparrow retina and the periphery of the Acadian flycatcher retina were examined for negative confirmation because the novel structure we sought is only present near the center of the Acadian flycatcher retina. Ultrathin cross sections (50 nm) were inspected using a JEOL-100CX electron microscope (JOEL Ltd., Tokyo, Japan).

Using ImageJ, we measured the length and width of 17 individual megamitochondria from MMOD-complexes and 28 mitochondria from traditional cones. We also measured the diameter of 152 small oil droplets from MMOD-complexes and 8 oil droplets from traditional cones.

### Retinal Wholemounting and Photoreceptor Distribution

One right eye from a least flycatcher, one right eye from an Acadian flycatcher, and two left eyes from Acadian flycatchers (4 total individuals) were used for retinal wholemeounting and photoreceptor distributions. Following euthanasia, eyes were immediately enucleated. We then hemisected the eye at the ora serrata with a razor blade and removed the vitreous humor using spring scissors, tweezers, and a dissecting microscope. We submerged the eyecup in cold 0.1 M phosphate buffered saline (pH 7.4) and placed it in a −20°C freezer for three minutes to facilitate the retraction of pigmented epithelium from the photoreceptor layer. The retina was removed from the eyecup by separating the choroid from the sclera and severing the optic nerve. We then gently peeled the choroid away from the retina and any pigmented epithelium that remained attached to the retina to avoid damaging the photoreceptor layer. We transferred the retina directly to a glass coverslip with the photoreceptor layer facing the coverslip. We then flattened the retina onto the glass coverslip by making several radial cuts with spring scissors and unrolling the retinal edges with a small paintbrush. We placed a drop of super glue onto each corner of the coverslip, which was then flipped onto a microscope slide and sealed with clear nail polish. We also wholemounted one white-throated sparrow retina for a qualitative comparison.

Retinae were then immediately imaged using SRS (Systematic Random Sampling) StereoInvestigator software (MBF Bioscience, Williston, Vermont), an Olympus BX51 microscope, and an Olympus S97809 camera (Olympus Corporation, Tokyo, Japan). A bright field illuminated image and an epifluorescent-illuminated image were captured at 275 to 280 locations on each retina. A counting frame of 50 μm X 50 μm was used at each 579 ± 39 μm X 621 ± 16 μm location with a mean ± SE area sampling fraction (i.e., the proportion of each location that was occupied by the counting frame) of 0.0071 ± 0.0008. Our mean ± SE Schaeffer’s Coefficient of Error of 0.046 ± 0.002 is less than 0.10, indicating that our stereological sampling strategy was appropriate^34^. We identified and counted all oil droplets within each counting frame using ImageJ (https://imagej.nih.gov/ij/). In birds, each cone type is associated with a specific oil droplet type. Oil droplets are oil and carotenoid-filled organelles at the apical end of the cone inner segment that spectrally filter light before the light reaches the visual pigment in the outer segment where phototransduction occurs. Therefore, the densities of each photoreceptor type can be quantified by counting oil droplets. Traditionally, there are five types of cones each associated with its own type of oil droplet. Transparent (T-type) oil droplets are found in ultraviolet or violet sensitive single cones, colorless (C-type) oil droplets are found in short-wavelength sensitive single cones, yellow (Y-type) oil droplets are found in medium-wavelength sensitive single cones, red (R-type) oil droplets are found in long-wavelength sensitive single cones, and principal (P-type) oil droplets are found in the principal member of double cones. We identified each of the five traditional cone types follow the criteria established by Hart^3^. Briefly, R-type oil droplets appear red under bright field illumination, Y-type oil droplets appear yellow under bright field illumination, T-type oil droplets appear clear under bright field illumination but are absent under epifluorescent illumination, C-type oil droplets appear clear under bright field illumination but fluoresce bright white under epifluorescent illumination, and P-type oil droplets appear clear to yellowish-green under bright field illumination but fluoresce dull white to dull yellow under epifluorescent illumination. In addition to the five traditional oil droplet types, we were easily able to identify the new MMOD-complexes as the obvious, large, orange triangles juxtaposed with the smaller, circular traditional oil droplets.

To visualize the distribution of photoreceptors across the retina, we constructed topographic maps in R (version 3.3.0) following Garza-Gisholt et al.^35^. We also calculated cell density ratios to determine if MMOD-complex photoreceptors are a subset of another type of photoreceptor type or a present at the expense of any single photoreceptor type. First of all, all five traditional oil droplet types are present in areas of *Empidonax* retinae containing MMOD-complex photoreceptors. To calculate cell density ratios, we took the number of oil droplets of a given type divided by the number of T-type oil droplets at each counting location. For example, the C-type ratio is calculated as the number of C-type oil droplets present in a single counting location divided by the number of T-type oil droplets at that location. And the Y-type ratio is calculated as the number of Y-type oil droplets present in a single counting location divided by the number of T-type oil droplets at that location, etc. We used T-type oil droplets as the denominator because they are the least abundant oil droplet type in the retina. Ratios were calculated for every counting location on each of the four *Empidonax* retinae. All the ratios from an individual retina were averaged to yield a single data point per individual. Those data were compared to typical retinae from Order Passeriformes – the same order that *Empidonax* flycatchers belong to – using the t.test function in R to determine if the cell ratios of the *Empidonax* retinae differed from expected values. Values for typical passerine retinae were garnered from Hart^3^ as the species-level means for six passerine species (*Entomyzon cyanotis, Manorina melanocephala, Parus caeruleus, Ptilonorhynchus violaceus, Sturnus vulgaris, Turdus merula*). Additionally, we used a one-tailed, paired t-test to determine if traditional cell density ratios were lower in MMOD-complex regions than regions lacking MMOD-complexes within *Empidonax* retinae, which would indicate that MMOD-complex photoreceptors are replacing other cones.

### Cross Sections

We performed histological cross sections on the right eye of one Acadian flycatcher and the left eye one least flycatcher. We enucleated, hemisected, and removed the vitreous humor of each eye as described above. Then we fixed the retinae in the eyecup with 4% paraformaldehyde in 0.1 M phosphate buffered saline (pH 7.4) for >24 hours. The eyecup was then cut into a 2 mm wide strip along the naso-temporal axis with the fovea in the center of the strip. We embedded the tissue in paraffin wax and serial sectioned the tissue at 5 μm using a Thermo Scientific Shandon Finesse ME microtome (Waltham, Massachusetts). Sections were stained with haemotoxylin/eosin in a Thermo Scientific Shandon Varistain 24-3.

We noted an unusual and abrupt change in thickness of the retinal tissue approximately midway between the fovea and the ora serrata in both the nasal and temporal directions. We measured the naso-temporal width of the thicker, central portion of the retina using a Zeiss Axio Imager.M2 microscope (Carl Zeiss Microscopy, Göttingen, Germany) and a Zeiss AxioCam MRm camera (Carl Zeiss Microscopy, Göttingen, Germany). To compare the width of the thick, central region to the width of the MMOD-complex area, we plotted counting locations from the previous section that contained MMOD-complex photoreceptors onto traces of the retinal wholemounts. We also used these plots in ImageJ to calculate the total percent of retinal area in which MMOD-complex photoreceptors were present.

### Microspectrophotometry

To determine the spectral absorbance characteristics of flycatcher oil droplets, the left eye of one Acadian flycatcher was analyzed using microspectrophotometry. Immediately after capture, the Acadian flycatcher was light deprived for three hours to facilitate visual pigment regeneration. Euthanasia, enucleation, hemisection, and vitreous extraction were all carried out under dim red light to avoid bleaching the photoreceptors. The retina was extracted from the eyecup in the same fashion as described above for retinal wholemounts, and we used a small paintbrush to brush away excessive pigmented epithelium that remained attached to the retina. We removed a ∼6 mm^2^ piece of the retina, placed the retina section on a glass slip, and macerated the retina section with a razor blade. The macerated retina was hydrated with a drop of phosphate buffered saline (pH 7.4) and sucrose water, cover slipped, and sealed with black nail polish to prevent desiccation.

We measured absorbance spectra of oil droplets and MMOD-complexes using a custom-built microspectrophotometer (Dr. Ellis Loew, Cornell University, Ithaca, NY; described in McFarland and Loew^36^). We used a dry Zeiss Ultrafluar Glyc objective (80x, NA 0.9) for viewing and a second objective (32x, NA 0.4) as a condenser. We viewed cellular structures using an EXVision Super Circuits CCD camera attached to a TFT Color LCD monitor that was covered in red Plexiglas. After identifying an isolated oil droplet or MMOD-complex, we took a baseline measurement in the surrounding interstitial space. Then, we measured the absorbance of the target structure in 1 nm increments from 350-750 nm. The absorbance of MMOD-complexes was measured through the thicker basal end and the thinner apical end of the structure separately. We attempted to bleach a MMOD-complex by exposing it to white light for 60 s to determine whether it behaved as an oil droplet or a visual pigment. Visual pigments bleach (i.e., stop absorbing light) under white light^37^, whereas oil droplets do not bleach.

Oil droplets are typically characterized by three parameters: λ_cut_ – the wavelength of maximal absorbance; λ_mid_ – the wavelength with 50% of maximal absorbance; and λ_0_ – the wavelength of 63% maximal absorbance. We determined these three parameters using the MATLAB program, OilDropSpec. OilDropSpec normalized the long wavelength arm of each oil droplet spectrum to one, found the wavelength where absorbance was 0.5 (λ_mid_), fit a trend line to the absorbance data 10 nm to either side of λ_mid_, determined the intercept, slope, and R^2^ of the trend line, used the trend line to determine the wavelength where the absorbance was 1 (λ_cut_), and calculated λ_0_. In total, absorbance spectra were measured for 1 T-type oil droplet, 1 C-type oil droplets, 17 Y-type oil droplets, 9 R-type oil droplets, 13 P1-type oil droplets, 4 P2-type oil droplets (there are two standard variations of P-type oil droplets), and 23 MMOD-complexes.

## Acknowledgements

Thanks to J. Lucas and A. Gleichsner for comments on previous versions of the manuscript. Funding for this project was provided by the National Science Foundation (IOS #1146986 to EFJ).

## Author Contributions

LPT discovered the MMOD-complex. LPT and EFJ designed the study. LPT, LBCT, RDD, BAM, PB, and DP carried out experiments and analyzed results. LPT and EFJ wrote the manuscript. All authors approved of its submission.

## Data Availability

All data generated during this study are included in this published article, its Supplementary Information files, or are available from the corresponding author.

