## Supplementary Materials for "A novel cellular structure in the photoreceptors of insectivorous birds"

**Supplementary Table S1 | Microspectrophotometry of *Empidonax* flycatcher oil droplets**

*No values are present for T-type oil droplets because they are transparent with no absorbance curve.

| | T-type* | C-type | Y-type | R-type | P1-type | P2-type | | MMOD-complex |
|---|---|---|---|---|---|---|---|---|
| | | | | | | a | b | |
| Mean $\lambda_{cut}$ (nm) | — | — | 500 ± 1.83 | 560 ± 2.38 | 433 ± 2.48 | 449 ± 3.68 | 485 ± 1.26 | 543 ± 0.92 |
| Mean $\lambda_0$ (nm) | — | — | 512 ± 2.18 | 575 ± 1.86 | 451 ± 2.84 | 457 ± 2.83 | 495 ± 1.25 | 562 ± 0.91 |
| Mean $\lambda_{mid}$ (nm) | — | — | 517 ± 2.31 | 581 ± 1.90 | 457 ± 3.11 | 459 ± 2.54 | 499 ± 1.26 | 569 ± 0.92 |
| Mean b | — | — | 0.088 ± 0.003 | 0.071 ± 0.005 | 0.065 ± 0.005 | 0.140 ± 0.014 | 0.105 ± 0.002 | 0.056 ± 0.001 |
| Mean $B_{mid}$ | — | — | 0.030 ± 0.001 | 0.024 ± 0.002 | 0.022 ± 0.002 | 0.048 ± 0.005 | 0.036 ± 0.001 | 0.019 ± 0.000 |
| $\lambda_{cut}$ of mean absorbance spectrum (nm) | — | 405 | 498 | 562 | 435 | 449 | 485 | 541 |
| $\lambda_0$ of mean absorbance spectrum (nm) | — | 412 | 511 | 575 | 447 | 456 | 495 | 559 |
| $\lambda_{mid}$ of mean absorbance spectrum (nm) | — | 415 | 516 | 580 | 451 | 459 | 498 | 565 |
| b of mean absorbance spectrum | — | 0.150 | 0.080 | 0.082 | 0.090 | 0.139 | 0.114 | 0.060 |
| $B_{mid}$ of mean absorbance spectrum | — | 0.052 | 0.028 | 0.028 | 0.031 | 0.048 | 0.039 | 0.021 |
| n | 1 | 1 | 16 | 8 | 13 | 4 | — | 23 |

**Supplementary Table S2 | Statistics on oil droplet density ratios**

Mean ± SE values can be found in Fig. 3a.

| Comparison | Ratio | t-statistic | Degrees of freedom | p-value |
|---|---|---|---|---|
| Typical passerine versus *Empidonax* | C-type:T-type | 0.33 | 8 | 0.752 |
| | Y-type:T-type | 0.87 | 8 | 0.411 |
| | R-type:T-type | 0.20 | 8 | 0.843 |
| | P-type:T-type | 0.16 | 8 | 0.876 |
| | MMOD-complex:T-type | 9.64 | 8 | **< 0.001** |
| *Empidonax* periphery versus *Empidonax* center | C-type:T-type | 0.67 | 3 | 0.725 |
| | Y-type:T-type | 3.61 | 3 | 0.982 |
| | R-type:T-type | 0.68 | 3 | 0.274 |
| | P-type:T-type | 1.21 | 3 | 0.059 |
| | MMOD-complex:T-type | 22.88 | 3 | **< 0.001** |

**Supplementary Table S3.** List of species used to construct a phylogeny of birds for which photoreceptor and oil droplet types are known. For statistical analyses, we only used species for which eye size and foraging data were also published; these species are denoted in the Used in Analysis column.

| Family | Species | Source |
|---|---|---|
| Dromaiidae | *Dromaius novaehollandiae* | Hart et al. 2016 |
| Struthionidae | *Struthio camelus* | Wright & Bowmaker 2001 |
| Rheidae | *Rhea americana* | Wright & Bowmaker 2001 |
| Anatidae | *Anas penelope* | Hart 2001 |
| Anatidae | *Aythya affinis* | Hart 2001 |
| Laridae | *Larus novaehollandiae* | Hart 2001 |
| Laridae | *Anous minutus* | Hart 2001 |
| Columbidae | *Stigmatopelia chinensis* | Hart 2001 |
| Columbidae | *Columba livia* | Bowmaker et al. 1997 |
| Alcedinidae | *Todiramphus sanctus* | Hart 2001 |
| Cuculidae | *Eudynamys scolopaceus* | Hart 2001 |
| Phasianidae | *Pavo cristatus* | Hart 2001 |
| Phasianidae | *Gallus gallus* | Hart et al. 2006 |
| Rallidae | *Gallunula tenebrosa* | Hart 2001 |
| Phalacrocoracidae | *Phalacrocorax varius* | Hart 2001 |
| Procellariidae | *Puffinus pacificus* | Hart 2001 |
| Accipitridae | *Aquila audax* | Reymond 1985 |
| Accipitridae | *Buteo buteo* | Mitkus et al. 2017 |
| Accipitridae | *Pernis apivorus* | Mitkus et al. 2017 |
| Accipitridae | *Milvus milvus* | Mitkus et al. 2017 |
| Accipitridae | *Accipiter nisus* | Mitkus et al. 2017 |
| Falconidae | *Falco berigora* | Reymond 1987 |
| Falconidae | *Falco peregrinus* | Mitkus et al. 2017 |
| Cacatuidae | *Cacatua roseicapilla* | Hart 2001 |
| Psittaculidae | *Melopsittacus undulatus* | Knott et al. 2012 |
| Psittaculidae | *Platycercus eximius* | Hart 2001 |
| Psittaculidae | *Trichoglossus chlorolepidotus* | Hart 2001 |
| Fringillidae | *Carduelis tristis* | Baumhardt et al. 2014 |
| Turdidae | *Turdus migratorius* | Unpublished data |
| Turdidae | *Turdus merula* | Hart 2001 |
| Turdidae | *Sialia sialis* | Unpublished data |
| Paridae | *Parus caeruleus* | Hart 2001 |
| Meliphagidae | *Entomyzon cyanotis* | Hart 2001 |
| Meliphagidae | *Manorina melanocephala* | Hart 2001 |
| Icteridae | *Molothrus ater* | Fernandez-Juricic et al. 2013 |
| Icteridae | *Agelaius phoeniceus* | Unpublished data |
| Icteridae | *Sturnella magna* | Tyrrell et al. 2013 |
| Sturnidae | *Sturnus vulgaris* | Hart 2001 |
| Passerellidae | *Zonotrichia albicollis* | This study |
| Passerellidae | *Spizella pusilla* | Unpublished data |
| Passeridae | *Passer domesticus* | This study |
| Ptilonorhynchidae | *Ptilonorhynchus violaceus* | Hart 2001 |
| Cardinalidae | *Cardinalis cardinalis* | Unpublished data |
| Corvidae | *Cyanocitta cristata* | Unpublished data |

| | | |
|---|---|---|
| Mimidae | *Dumetella carolinensis* | Unpublished data |
| Sittidae | *Sitta carolinensis* | Unpublished data |
| Tyrannidae | *Empidonax virescens* | This study |
| Tyrannidae | *Empidonax virescens* | This study |

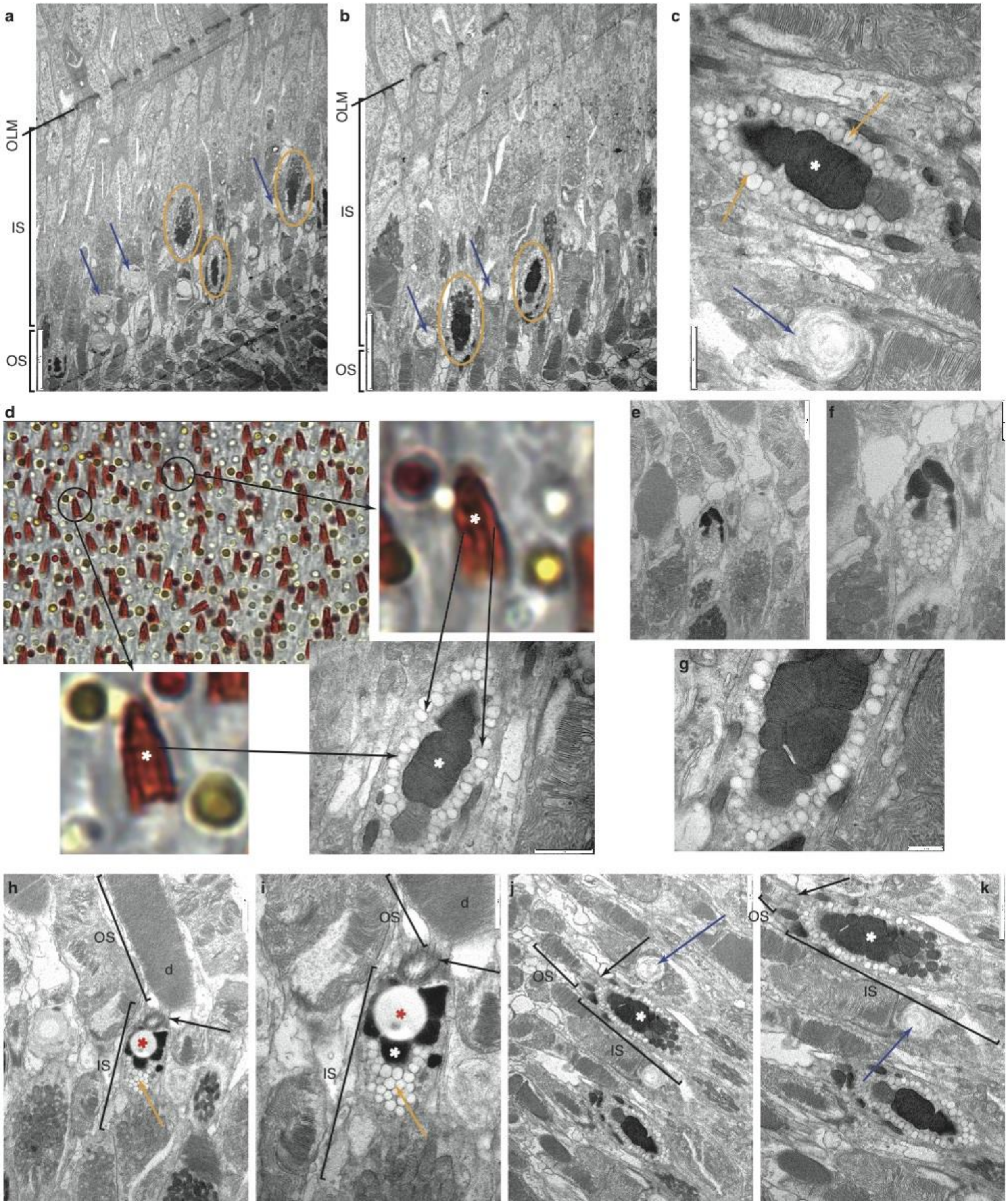


**Supplementary Figure S1 | Images of MMOD-complex photoreceptors.** (a-c, e-k)

Transmission electron microscopy of the MMOD-complex photoreceptors in *Empidonax* flycatchers. (d) The shape, arrangement, and location of the MMOD-complex observed under transmission electron microscopy correlates with that of the wholemount under light microscopy. OLM refers to the outer limiting membrane, IS refers to the inner segments, OS refers to the outer segments, and *d* refers to photoreceptor discs. Orange ellipses encompass MMOD-complexes, orange arrows indicate the small oil droplets surrounding megamitochondria that confer the orange color on the wholemount, blue arrows indicate traditional oil droplets, white asterisks indicate electron-dense megamitochondria cores, and red asterisks indicate artifacts. Black arrows in (h-k) indicate connecting cilium between inner and outer segments. The MMOD-complexes shown were only present in the two samples form the center of an *Empidonax* retina. No similar structures were present in the peripheral *Empidonax* sample or any of the house sparrow samples.

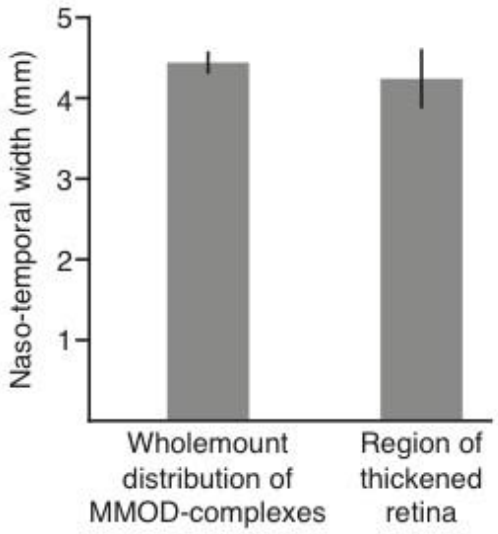


**Supplementary Figure S2 | Retinal coverage of MMOD-complex photoreceptors.** The distribution of MMOD-complex photoreceptors as measured on a retinal wholemount (image in Fig. 2j) corresponds to the thickened retinal region as measured on a cross section (image in Fig. 2k). This suggests that the MMOD-complex photoreceptors increase the demand for cells in other retinal layers, leading to a substantial thickening of the retinal tissue in regions where MMOD-complex photoreceptors are present. The MMOD-complex region covers the central $27.4 \pm 1.7$ $mm^2$ or $24.6 \pm 0.9\%$ of the *Empidonax* retina.
